# The ultrafast snap of a finger is mediated by skin friction

**DOI:** 10.1101/2021.08.25.457533

**Authors:** Raghav Acharya, Elio J. Challita, Mark Ilton, M. Saad Bhamla

## Abstract

The snap of a finger has been used as a form of communication and music for millennia across human cultures. However, a systematic analysis of the dynamics of this rapid motion has not yet been performed. Using high-speed imaging and force sensors, we analyze the dynamics of the finger snap. Our analysis reveals the central role of skin friction in mediating the snap dynamics by acting as a latch to control the resulting high velocities. We evaluate the role of this frictional latch experimentally, by covering the thumb and middle finger with different materials to produce different friction coefficients and varying compressibility. In doing so, we reveal that the compressible, frictional latch of the finger pads likely operate in a regime optimally tuned for both friction and compression. We also develop a soft, compressible friction-based latch-mediated spring actuated (LaMSA) model to further elucidate the key role of friction and how it interacts with a compressible latch. Our mathematical model reveals that friction plays a dual role in the finger snap, both aiding in force loading and energy storage while hindering energy release. Our work reveals how friction between surfaces can be harnessed as a tunable latch system and provide design insight towards the frictional complexity in many robotics and ultra-fast energy-release structures.

## 1. Introduction

### (a) History of the Snap

The earliest recorded representation of the finger snap dates back to as early as 320 BCE in Greece, where a piece of pottery depicts Pan, the god of the wild, dancing with his right hand curled in the position of a finger snap (Fig. 1A,B). Furthermore, the finger snap was commonly used by the Ancient Greeks to keep rhythm [1]. However, the finger snap is not confined to just one culture. Its simplicity and effectiveness at creating a sharp sound has been incorporated into many cultures, including the Flamenco dances of Spain, in which it is referred to as “chasquidos” [2]. The finger snap has even made its way into popular media, featuring prominently in 1961 film West Side Story and in Avenger movies (2018, 2019). Today, the finger snap has widespread applications, as gestures of greeting in Liberia (similar to a handshake) [3], as applause at poetry readings (instead of clapping), for simultaneous localization of multiple microphones [4], as a form of echolocation used by blind people [5], as part of psychological research into auditory development [6], or for biometric authentication of digital devices [7].

**Figure 1.**
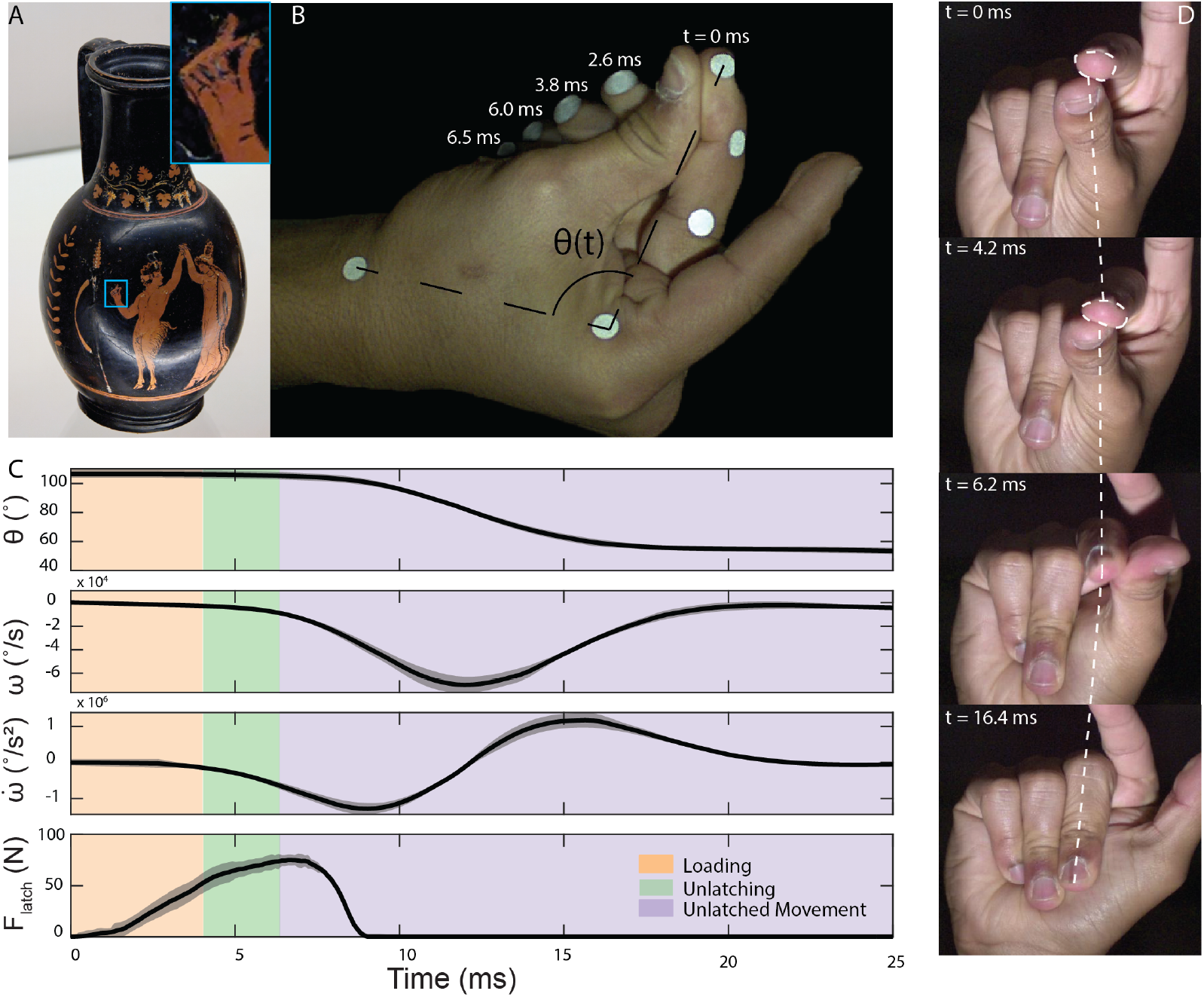
The finger snap is a three-phase, predominantly 1-D motion exhibiting high speeds and accelerations. **A**. A piece of pottery from 320-310 B.C.E depicting Pan, the Greek god of the wild, dancing with a Manead with the hand curled in the shape of a finger snap. Images are public domain from [22]. **B**. Composite image of the motion at different timestamps of the snap from a side view. **C**. Kinematics and dynamics of the finger snap (n = 5). Angle measurements taken between points on wrist, knuckle, and tip of finger. Force measurements taken via tactile pressure sensor placed between middle finger and thumb during snap, aligned such that force reading reaches 0 at peak acceleration. **D**. Stills of a finger snap from the front showing the visible compression of the fingertip as energy is stored before being released and causing the nearly 1-D motion of both the thumb and the middle finger.

### (b) Finger snaps as a LaMSA System

The motion we describe as a snap, referring in general to the contact of two appendages together as force is built and energy is stored before a rapid release and motion of one or both appendage, has been observed in numerous organisms. Multiple termite species, including *Termes panamaensis* [8] and *Pericapritermes nitobei* [9], as well as a species of ant known as *Mystrium camillae* [10], have all been observed pressing their mandibles together to generate ultra fast motion in a manner reminiscent of the finger snap that humans perform. Although the snapping behavior of various biological organisms differs in terms of purpose and function, the mechanism behind snapping may be classified as a Latch Mediated Spring Actuated (LaMSA) system [11]. A LaMSA system is one where energy is loaded in a mass-spring system by an external motor over a relatively long period of time before being held in place with a latch. Ultrafast movement is achieved when the latch is rapidly released, allowing the stored potential energy to explosively launch the mass in a relatively short period of time. Many biological organisms exploit this principle using biological springs and latches to achieve various functionalities. Some of these organisms include trap jaw ants, froghoppers, mantis shrimps, and the aforementioned snapping ant and termite species [12, 13, 14]. While the roles of the latch geometry and spring structures in snap-based LaMSA systems have been explored [11, 14, 15, 16], one key aspect of snapping systems that has yet to be explored in detail is that of friction. Friction has been hypothesized to play a key role in ensuring successful loading and unlatching of LaMSA systems [17] but it has not been analyzed.

In the case of the finger snap, we hypothesize that the arm muscles act as a motor to load potential energy in the tendons of the fingers and arms, which act as springs (Fig. 1B) [18, 19, 20]. The skin friction between the middle finger and thumb assists in the latching of the middle finger but also hinders unlatching and motion, playing a dual role in the dynamics of the snap. We begin analysis of the role of friction by experimentally varying the friction coefficient and compressibility of the materials covering the skin. We then develop a mathematical model that incorporates friction with a LaMSA system that can qualitatively capture the trends observed experimentally. Using this model, we reveal the role that friction plays in mediating the finger snap.

## 2. Methods

### (a) Kinematic Analysis

To measure and analyze the kinematics of a finger snap, we use 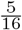 inch diameter circles of reflective tape and place them on the wrist joint, the base of the fingers (metacarpophalangeal joint), the first knuckle (proximal interphalangeal joint) of the middle finger, the second knuckle (distal interphalangeal joint) of the middle finger, and the tip of the middle finger of the snapper (Fig. 1B). We record all snaps using a Chronos 1.4 high speed camera at 4082 fps using a Nikon AF-S DX Nikkor 35 mm lens. Different surfaces (latex Rubber, Nitrile, Lubricated Nitrile) are placed over the middle finger and thumb to adjust friction and determine their effect on the snap. The lubricant used is water-based lubricant (Cetaphil moisturizer). Five snaps from three different people for each surface are analyzed. A new nitrile glove is used for each trial to prevent material fatigue, and fresh lubricant is applied before each trial for repeatability. The nitrile glove that best suited the hand size of the snapper was used. To prevent muscle and joint fatigue, the person performing the finger snap rests for one minute between snaps. See SI Fig S4 for data and variability from different individuals.

In the case of the nitrile-covered thimble snap, copper thimbles were placed on the middle finger and thumb. To secure the thimble and to maintain a standard friction coefficient, a nitrile glove is worn over the thimble. Five snaps from one person are recorded and analyzed the results of which are shown in Figures 2 and 3.

**Figure 2.**
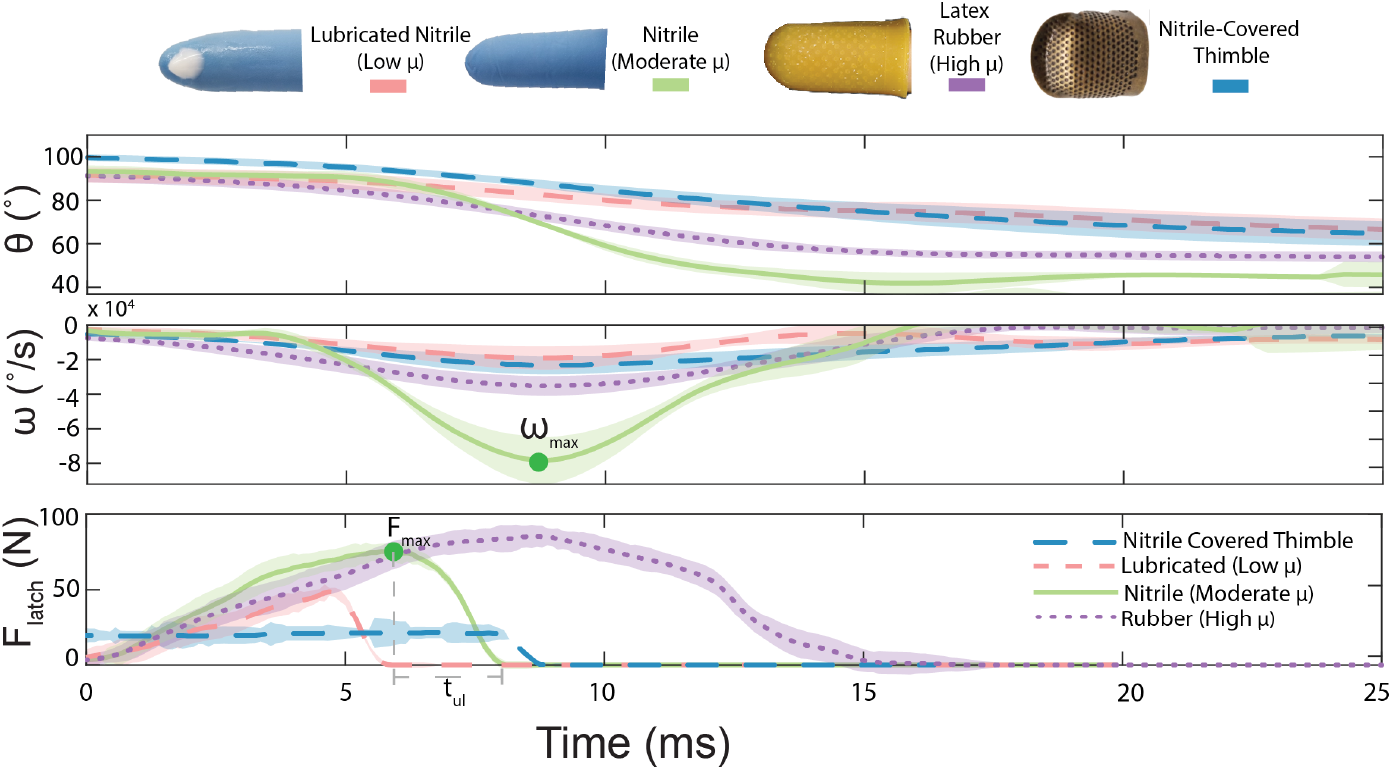
Effect of friction and compression on the finger snap. Fingers are covered in lubricated nitrile (low *µ*, pink), nitrile (moderate *µ*, green), latex rubber (high *µ*, purple), and a nitrile covered thimble (low contact area, blue) and angular displacement, velocity, and normal forces are reported. For lubricated nitrile, nitrile, and latex rubber experiments, N = 5 snaps are analyzed from three different people. For the nitrile covered thimble, five snaps are analyzed from one person. Snaps performed with lubricated nitrile and nitrile-covered thimble on fingers required longer than the shown 25 ms for the middle finger to reach a resting angle and this return to resting angle is omitted from the graph. The shaded areas represent variance of measurement at each point in time.

**Figure 3.**
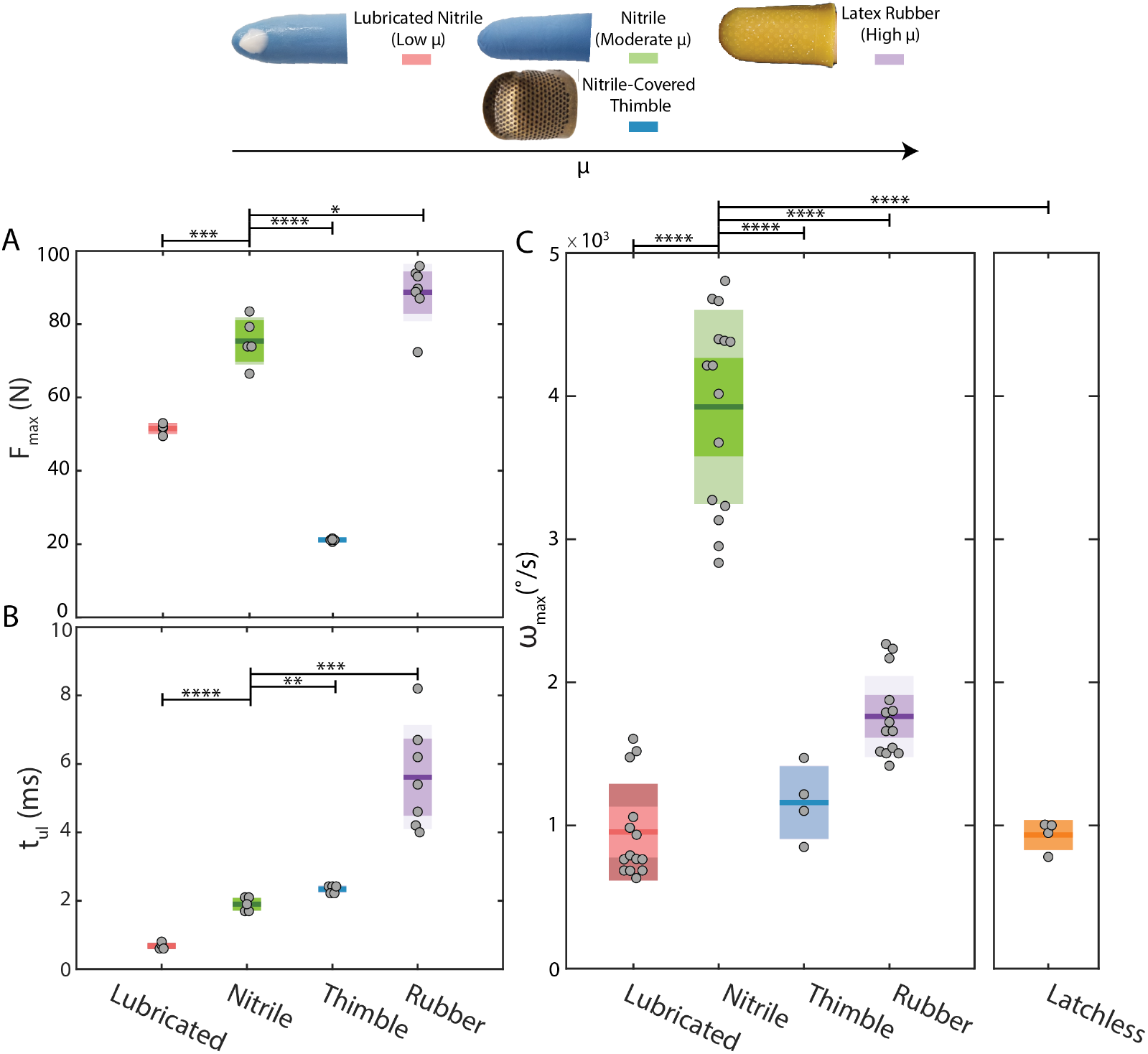
A moderate surface friction between thumb and finger results in highest observable velocities and accelerations. **A**. Variation of *F*_*max*_ with varying frictional surfaces. *F*_*max*_ represents the amount of force stored in the physiological springs within the finger system. The results show that the greater the *µ*, the larger the force that can be stored in the aforementioned spring. **B**. Variation of *t*_*ul*_ with varying frictional surfaces. *t*_*ul*_ is defined as the time of contact between the two fingers from first motion and therefore serves as a good indicator of how much energy is lost due to friction. It can be seen that *t*_*ul*_ increases with increasing *µ* indicating that as the friction increases, more energy is lost. **C**. Variation of *ω*_*max*_ with varying frictional surfaces. It can be observed that *ω*_*max*_ is highest for the nitrile covered snap which has a moderate coefficient of static friction compared to a lubricated covered snap (low *µ*) and the latex rubber covered snap (high *µ*). In addition, nitrile covered snap is orders of magnitude higher than a “latchless” snap, where no resistance by the thumb is given to the middle finger and pure muscle motion is allowed. In this example, it is also notable that the *ω*_*max*_ of the lubricated snap is very similar to that of the latchless snap, indicating that the low *µ* has disrupted the motion of the snap drastically.

A custom MATLAB script is used to track each dot during its motion using k-means clustering to locate the centroid of each dot, returning x-y positional data in pixels. The angular position is quantified by calculating the angle between the dots on the wrist, the base of the finger, and the tip of the finger. The angular velocity and acceleration are then calculated as the first and second order derivatives of the angular motion data after being smoothed using a zero order, 17 frame Savitzky-Golay filter. While the distance between the tip of the finger and the base of the finger changes slightly as the joints moved, we found this change to be small enough to not affect analysis. Pairwise t-tests are performed on the maximum velocities between nitrile and each surface tested to determine significance of difference.

### (b) Measuring Coefficients of Friction

A 100 mm linear actuator (Nema 23 Stepper Motor, CBX1605-100A) driven by a one axis controller stepper motor driver board (HiLetgo, TB6560 3A) is mounted horizontally on a sheet of acrylic with a micro load cell (CZL639HD, Phidgets Inc) on the moving arm (Fig. SI 2A). The actuator is controlled by an Arduino through a load cell amplifier (HX711 IC, Sparkfun). The load cell is connected to an Arduino board converting input voltage data to force values. The load cell is calibrated by placing attaching a known weight to the load cell with a string and allowing the weight to hang under the influence of gravity such that the tension on the string is horizontal. The voltages measured are used to build a calibration curve. A schematic of this experimental set up can be found in SI Fig 2A.

To measure the frictional coefficients of various materials, samples of the materials are affixed to an acryllic sheet and the bottom of a petri dish. Known weight is added to the petri dish, which is then attached to the string which connects to the load cell. The linear actuator is activated to move at a fixed velocity. The load cell readings are recorded and analyzed to find the steady state reading, which is then used to determine the coefficient of friction for the two materials. Five trials are performed for each material (nitrile, lubricated nitrile, latex rubber). Example plots and the experimental results can be found in SI Fig 2B and 2C respectively.

### (b) Dynamic Force Analysis of the Finger Snap

To measure the force developed between the fingers during the snap, we place an FSR Interlink 402 resisitive force sensor between the thumb and middle finger (SI Fig 2). This FSR has a diameter of 14 mm, which is comparable to the approximately 14.5 mm and 15.5 mm diameters of the elliptically shaped contact area of the finger pad. These diameters are measured by pressing the thumb and middle finger with high force against a clear sheet of glass and measuring the diameters of the elliptical contact area. The FSR is connected to a voltage divider and sends analog voltage data to an Arduino board running a script which converts the input voltage data to force values. After calibrating the voltage dividers for the FSR using manufacturer data, we validate the results by placing weights of known masses (3 kg to 10 kg) on the FSR.

Force data is collected for N = 10 snaps made while wearing a nitrile glove, changing the glove between trials and allowing at least one minute between snaps to prevent fatigue. Five of these snaps are performed to be as fast as possible, and an audible snap was detected. The other five are intentionally performed to be “weak” such that no sound was detectable. The differences between these snaps is further detailed in SI Fig 5, which reveals how the fastest snaps for a given set of experimental conditions can be distinguished from “weaker” snaps by the forces achieved and the variation in unlatch times. These differences are used to identify outliers in force data taken for different sets of conditions for further examination.

Force data is collected for 5 snaps made while wearing a nitrile glove with lubrication, while wearing latex rubber on both fingers, and while wearing a metal thimble on both fingers underneath the nitrile glove. In this thimble case, to prevent the issue of partial loading, 13 mm diameter circles of 1 mm thick acrylic were cut and used to ensure complete compression of the sensor while the snap was performed. The force data from all trials is then analyzed for two key features - the maximum normal force (*F*_*max*_) applied by the finger, and the unlatch time (*t*_*ul*_) (Fig 2). Unlatch time is defined as the time from when the maximum normal force is reached till zero force is read, which signifies a complete unlatching of the middle finger and thumb. Pairwise t-tests are performed on both *t*_*ul*_ and *F*_*max*_ between nitrile and each surface analyzed to determine significance. These results can be found in Figures 2 and 3 and SI Table 3.

### (c) Mathematical Modelling

This model is developed based on experimental observations and a more detailed explanation of the model development can be found in section 3C. A model schematic is illustrated in Fig 4B. It consists of a load of mass *m*_*l*_ (representing the middle finger) mounted on a spring with stiffness *k* (representing the elastic structures of the forearm and hands) displaced to equilibrium length *y*_*eq*_ by a linear force-velocity loading motor (representing the muscles of the forearm) with a maximum motor force. The load is held in place by a circular latch of radius *R* and mass *m*_*l*_ (representing the thumb). If the *y*_*eq*_ is small enough that the force of friction can oppose the motion of the load and latch, then it will produce a stable system. Once the spring has been loaded, the latch accelerates by a linear force-velocity [21] unlatching motor (representing the muscles which move the thumb) with a defined maximum force and velocity.

**Figure 4.**
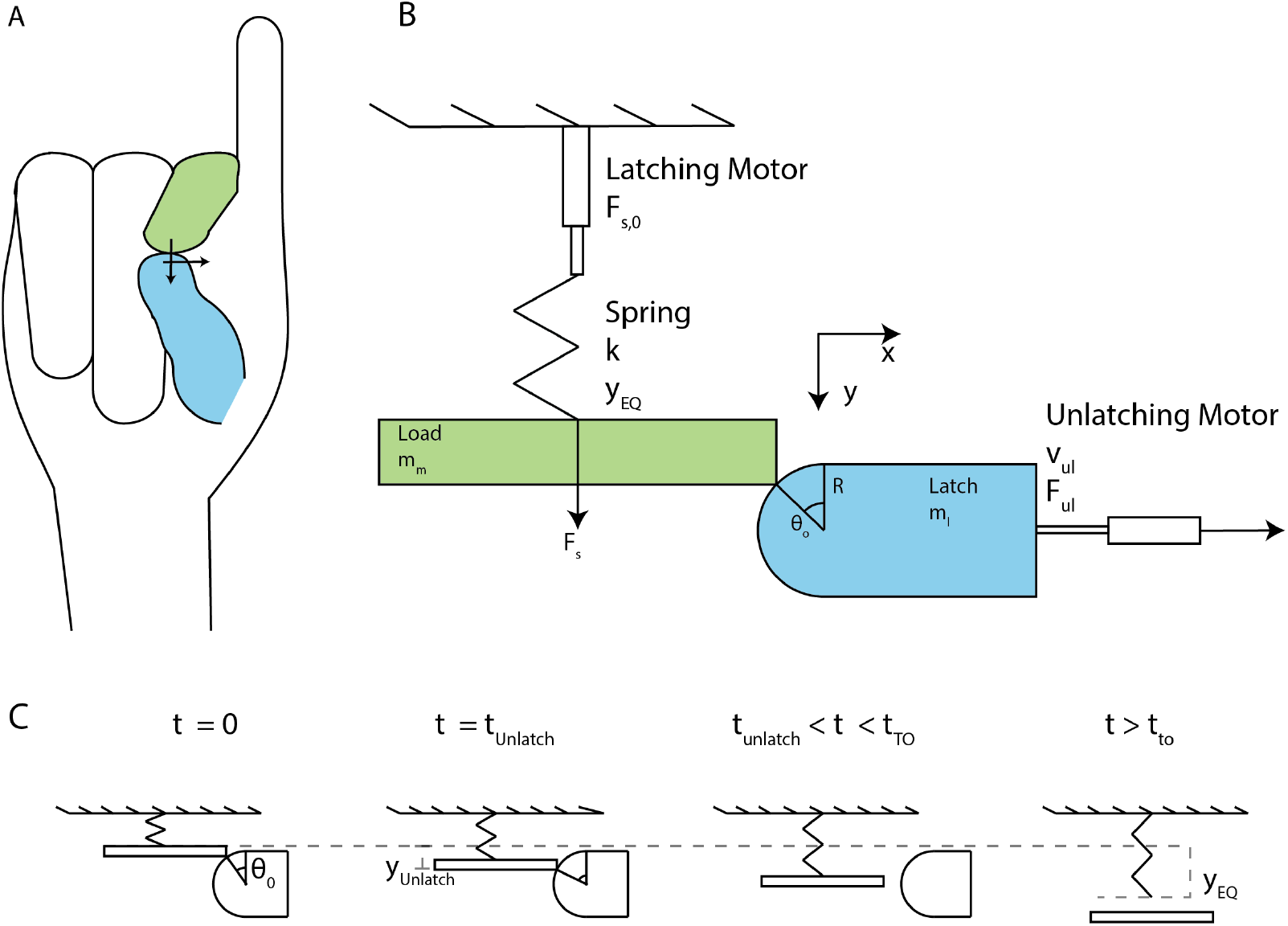
Finger snap modelled as a 1-D latch spring system. **A**. Schematic of finger snap showing motion of thumb (blue) which acts as the latch and middle finger (green) which acts as the load. **B**. Analogy to traditional latch mediated spring actuated systems where a latch (blue) allows for the storage of energy in the spring which later drives a load mass (green) as it unlatches. **C**. These schematics show the evolution of the system over time. Initially the system begins with the spring compressed and the load and latch positioned at angle *θ*_0_. The system moves as the unlatching motor acts on the latch, causing the load to accelerate in the positive y direction until the unlatch time is reached, which is the last time the latch and load are in contact. After this point, the load continues to accelerate solely due to the spring force with no other forces acting on it. This continues until the take off time is reached, which occurs when the equilibrium spring position is reached. After this point, the load continues to move without any external force.

The latch and load have a coefficient of friction *µ* and start with the corner of the load mass offset by an initial angle *θ*_0_ with respect to the center of the latch. The model checks how much energy can be loaded in the spring based on the *µ* and the *θ*_0_ before displacing the spring to *y*_*eq*_. Then, the unlatching motor begins to apply force to the latch, accelerating the load. Equations 3.1, 3.5, 3.4, and 3.7 are used with an ordinary differential equation solver to determine the position, velocity, accelerations, and forces of the load mass and latch at each time step. The unlatching point is defined as the time when the normal force between latch and load goes to zero. After this point, the latch continues accelerating solely due to the unlatching force while the load continues accelerating solely under the effect of the unloading of the remaining elastic potential energy within the spring. Once the spring reaches equilibrium, the load is travelling at its fastest, and is assumed to decouple from the spring instantaneously. This point is accordingly defined as the take-off time and after this point, the load continues at the constant take-off velocity with no other forces acting upon it.

The mathematical model was coded in Matlab R2020b. All functions and scripts used can be found at this link: https://github.com/bhamla-lab/FingerSnap_2021.

## 3. Results and Discussion

### (a) Kinematics of the finger snap

How fast is a finger snap? We use high speed imaging at 4082 fps to quantify the kinematics of the moving fingers (see Methods and SI Movie 1). We divide the finger snap into three different phases: loading, unlatching and unlatching movement (Fig. 1C). In the loading phase, normal force is first built up by pressing and compressing the thumb and middle fingers together (Fig. 1D). During this phase, energy is stored in the deformation of elastic components of the hand and forearm. We hypothesize these spring-like elements are the tendons in the fingers or forearm. [18, 19, 23]. However, due to the complex structure of a human arm, the exact spring source remains an open area of inquiry. We hypothesize that by preventing motion, the friction between the two fingers plays a key element by acting as a ‘latch’. Eventually, the unlatching process begins with the thumb moving laterally and the middle finger quickly sliding past the thumb, thus starting the snap motion. During the snapping motion, the finger rapidly rotates by an angle *θ* = 53.5 *±* 6.7*°* in six milliseconds, *t* = 7.1 *±* 3.4 ms (N=5), nearly 30 times faster than the blink of an eye (Fig. 1C) [24, 25]. The finger continues its roughly 1-D motion (Fig. 1D) until it impacts the palm, reaching a *ω*max = 7.8 *±* 1.4 *×* 10^3*°*^*/s* and 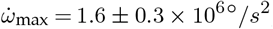, decelerating to a stop with *a*_dec_ = 1.8 *±* 0.4 *×* 106*°/s*2 (Fig. 1C). This impact generates weak shock waves (similar to a hand clap [26, 27]) that results in the characteristic pop sound.

To put these angular velocities and accelerations into context, the fastest rotational motion observed in humans has been in professional baseball pitches, with record-setting angular velocities of 9.2*×*10^3^ °*/*s and angular accelerations of 6.0*×*10^5^ °*/*s^2^ [28, 29]. Compared to these professional baseball pitch motions, recorded human finger snaps exhibit angular velocities within the same order of magnitude as the pitch, but exhibits angular accelerations almost one order of magnitude larger. Thus, to the best of our knowledge, the snap of a finger represents one of the fastest recorded rotational accelerations achieved by the human body. However, these velocities and accelerations are still low compared to those achieved by other snapping living systems such as in *Mystrium camillae*, the dracula ant, which can achieve velocities up to *v* = 2.4*×*108 °*/*s and accelerations up to *a* = 1.5*×*1016 °*/*s2 [10].

### (b) Friction Mediated Latch Dynamics

What role does the skin friction play in a snap? Unlike many spring-latch systems where unlatching is mediated by the latch geometry [15], finger snaps are unique in that they heavily mediated by friction between surfaces. We hypothesize that while the skin friction ultimately aids in the build-up of force during the energy storage phase by acting as a latch, it also hinders motion in the unlatching phase as it dissipates energy.

We explore this hypothesis by systematically changing the friction coefficient (*µ*) between the thumb and middle finger with different materials. We note that for creating a controlled, reproducible surface and eliminating variability due to skin sweat, we use nitrile gloves as our control case. We show that it faithfully replicates the performance of a bare skinned finger snap (SI Fig. S1). To better quantify the differences in coefficient of friction for each material, we experimentally measure it using the force set up described in Methods Section B and SI Fig 2.

Using a force sensor, we measure the temporal variation of the normal force between the middle finger and thumb during the loading phase as shown in Fig. 2. These force profiles reveal two features of interest that change with each surface tested: maximum normal force achieved before the snap occurs (*F*_max_) and the ‘unlatching’ time (*t*_ul_) defined as the time taken for the maximum normal force to decay to zero (Fig. 2). For a nitrile covered snap (control, nitrile on nitrile *µ* = 0.214 *±* 1.1 *×* 10^*−*4^, SI Fig 2), the *F*_max_ = (75.0*±*6.4) N and *t*_ul_ = (1.9*±*0.2) ms (Fig. 2).

We next lubricate the nitrile surface with a water based moisturizer thus reducing the coefficient of friction dramatically by a factor of 45X (low friction latch, lubricated nitrile on lubricated nitrile *µ* = 4.95 *×* 10^*−*3^ *±* 0.77 *×* 10^*−*3^, SI Fig 2, SI Movie S2) [30]. As expected, reducing *µ* results in both decreased force loading and unlatch times approximately by factors of 1.4 and 2.8. compared to a normal snap, respectively (Fig. 3). Thus despite still being able to compress, disrupting the latch through lubrication diminishes the ability of the fingers to controllably build-up adequate normal force and leads to reduced snap velocity, 4X lower than the nitrile-covered snap (Fig. 3).

To explore the effect of the compression seen in the finger (Fig. 1D) without altering the friction coefficient, snaps are next performed with a metallic thimble on both the thumb and middle finger below the nitrile glove. Due to the incompressible surface, the force loading decreased by a factor of approximately 3.8X (SI Movie S) while the unlatch time increased slightly by a factor of 1.2X (Fig. 3A,B). These changes in force dynamics resulted in a drastic decrease in snap velocity, approximately 3.4X lower than the normal nitrile covered snap (Fig. 3C). These results demonstrate the importance of the compressibility of human skin and the subsequent increase in friction as a result of increased contact area in the force loading. This hypothesis is discussed in greater detail in the Modeling of Finger Snaps as a Compressible Skin-Friction Latch section.

We also explore the effect of enhancing the friction coefficient to create a high friction latch (SI Movie S3). Using a latex rubber cover, *µ* of the frictional latch increases by a factor of approximately 5.8X (latex rubber on latex rubber *µ* = 1.24 *±* 0.14, SI Fig 2) without affecting the compressibility of the two surfaces. These latex rubber covers increase normal force between fingers by a factor of approximately 1.2X compared to the regular snap (Fig. 3A). Counter-intuitively, the large loading force does not necessarily lead to improved snap velocity. When compared to the control snap, the finger shows a decrease in angular velocity by 2.2X (Fig. 3C). This decrease in velocity is due to the significant dissipation of energy during unlatching of the high friction surfaces and is reflected in the relatively long time it takes for the fingers to unlatch and freely rotate, with *t*_*ul*_ for latex rubber being nearly 2.9X greater than *t*_*ul*_ for nitrile (Fig. 3B).

All these systems show higher angular velocities than a case where motion is driven without the presence of the latch, where the middle finger is simply moved as fast as possible on to the palm without opposition from the thumb (SI Movie S4). In this ‘latchless’ case, we observe the lowest velocity, revealing that the frictional latch plays an important role in storage and rapid release of energy for this high-speed motion (Fig. 3A).

Altogether, the highest velocity finger snaps are observed when the fingers are covered in a moderate *µ* surface which permits the compression of skin. The results show that covering the fingers with lubricated nitrile (low *µ*), latex rubber (high *µ*), or a copper thimble (limiting compression) leads to significantly decreased *ω*_*max*_ 3. This suggests that the soft frictional latch may be operating in a regime that is optimally tuned in both friction and compression (Fig. 3A). High friction enables greater force loading (Fig. 3B), but also increases energy dissipation during unlatching as represented by the increase in *t*_*ul*_ (Fig. 3C). Low friction reduces energy dissipation from the latch (making it an energy efficient latch), but limits the amount of normal force that can be loaded in the system before becoming unstable and slipping. Additionally, low compression reduces the contact area between fingers, making the system more unstable due to a lower force of friction and therefore a dramatic reduction in force loading. From a kinematic standpoint, maximizing the velocity of the system requires not only a quick-release latch but also one that can enable build-up of maximum force before slipping. Therefore, to achieve the highest velocities with the system, these experimental results suggest the presence of an optimum friction and compressibility that both maximizes the potential force buildup between the fingers (*F*_*max*_) as well as minimizes the dissipated energy by lowering the unlatch time (*t*_ul_).

### (c) Modeling of Finger Snap as a Compressible Skin-Friction Latch

Inspired by previous mathematical models of LaMSA systems [15], we develop a simplified mathematical model of finger snapping in order to understand the role of friction in this soft body system qualitatively. By observing the trends this model predicts and qualitatively comparing them to those found in finger snap experiments, we will demonstrate that friction plays a key role in releasing the storing and releasing elastic energy. We will also use this model to explore the general features of the energetics involved in a friction-mediated LaMSA model.

The simplified model we develop builds off our previously published model of spring-driven movements described in detail in [15, 31]. This model uses a loading motor to compress a spring against a latch, storing an amount of elastic energy determined by both the force characteristics of the loading motor and the material and geometric properties of the spring. The latch has both geometric and frictional properties, which keeps the spring from prematurely releasing energy. Finally, an unlatching motor applieas force to the latch, moving it and allowing the spring to recoil, driving the load mass (Fig. 3). An important element of the model is the interaction between the latch and the recoiling system, which determines they energy dissipation of the system.

Each aspect of this model is analogous to some aspect of the finger snap. The loading motor is analogous to the muscles of the hand and forearm and the spring it compresses to store energy is likely analogous to the deformation of the tendons and fingers [18, 19, 23]. The latch is analogous to the thumb in the finger snap, which is accelerated away by the palm muscles, represented by the unlatching motor. The removal of the thumb allows the release of the energy stored in the deformed tendons or fingers (spring) which drives the load mass, representing the middle finger. The middle finger and thumb move approximately perpendicular to each other in 1-D. Therefore, the analogous components of the load mass and latch are also limited to 1-D motion perpendicular to each other. By linking components of this basic LaMSA model to the various parts of the finger snap, we begin to make additions to this base model to better capture the frictional trends of the system.

The first major addition to this model revolves around the loading motor (muscles), that stores energy in the deformation of the spring (deformation of tendons or fingers). Biological motors have a limit to the force they are capable of exerting [32]. In order to capture this aspect of the finger snap, we impose an artificial maximum on the force that can be exerted by the loading motor. This in turn establishes an upper limit on the amount of elastic energy that can be stored within the springs. For the purposes of these simulations, we set an approximate maximum value of *F*_*s*,0_ to 70 N which corresponds to maximum value observed in experiments (Fig. 3).

In previous LaMSA systems, the latch geometry has been assumed to be sufficient in preventing load release [21, 31, 33]. This assumption has been justified by limiting latch and load motion to 1-D motion and by positioning the latch and load during force loading in such a way that there would be no horizontal force vectors acting on the latch as a result of the load. In a circular latch, this requires precise positioning of the latch and load to have their point of contact be perfectly perpendicular to the direction of the spring force. Such a case has *θ*_0_ = 0, where *θ*_0_ is defined in Fig. 4B. In cases where *θ*_0_ *>* 0, where the load mass is positioned along the curve of the latch during loading, the latch geometry alone would not be sufficient to prevent motion, as the storage of force in the system would cause the load mass to push the latch away. In such cases, as the loading motor begins to store energy in the spring, the load mass will exert a force with a horizontal component on the latch, pushing it aside and negating the effectiveness of the latch. However, if there were sufficient friction between load and latch, the latch would not be pushed away and more force could be added.

Another useful way to consider the idea of *θ*_0_ is through the lens of instability. If the friction between the latch and load is high, then the maximum *θ*_0_ before slipping occurs is greater. Such a system is highly stable and as a result, more energy could be easily stored without slipping. This hypothesis is supported by experimental evidence, as the latex rubber-covered snap (high *µ*) allowed significantly greater force buildup than the nitrile-covered snap (moderate *µ*). The opposite is also true. If the friction between latch and load is low, then slipping occurs at much lower *θ*_0_, making the system much less stable and making it harder to store energy. This hypothesis is also supported, as the lubricated snaps (low *µ*) exhibited significantly lower stored forces compared to nitrile-covered snaps (moderate *µ*). These results show that friction plays a crucial role in allowing sufficient energy storage, likely by providing an additional force opposing and preventing the premature motion of the load mass. Therefore, to capture this behavior in the model, an initial *θ*_0_ is set for each system tested. As shown in Equation 3.2, *θ*_0_ determines how much energy can be loaded in the spring by a force balance between the normal force from the spring and the force of friction opposing motion.

The basic LaMSA model makes the assumption that both the load mass and latch are rigid bodies, allowing for the application of basic laws of intersurface friction. However, we have previously shown that the pads of fingers are soft bodies which deform significantly as finger snap begins (Fig. 1D), and that this compression results in a change in finger-finger contact area that impacts the force loading through the thimble tests (Fig. 3). Previous work on friction in soft bodies has shown that the change in area due to deformation can lead to more complex forms of friction [30, 34]. Thus, we use a simplified model used previously to model soft skin friction for finger pads [35, 36] which follows the form:

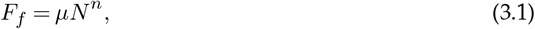

in which *F*_*f*_ represents the force of friction between two surfaces with relative coefficient of friction *µ* and normal force *N*. The constant *n* represents a constant that captures the nonlinearity of the frictional force that arises due to deformation and has been measured for soft bodies to be less than one (we use n=0.85, see the discussion below). In soft body systems such as finger pads, 0 *< n <* 1 which reflects the rapid initial increase in friction due to the increase in contact area [36]. However, past a certain force threshold, the area of contact does not further increase, resulting in a more linear relationship between friction and normal forces, which is also reflected by this empirical model. We acknowledge that this simple empirical formula for friction is a rather simplified method for modeling soft body friction. However, it does provide a first order approximation to the nonlinear behavior in two compressive touching bodies.

Because of the initial nonlinearity of this friction model caused by *n <* 1, the friction force grows slower than the normal force. Since the normal force is dependent on the spring force, this nonlinearity imposes a maximum spring force that can be reached for any given *µ* and *θ*_0_. Beyond this maximum spring force, the horizontal component of the normal force overcomes that of the friction force causing the latch to slip and the load to push the latch away. This relationship can be found by combining the equation for friction 3.1, with the following inequalities, derived from force balances on the latch and load, which must be true for energy storage to occur during latching:

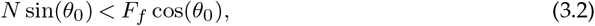

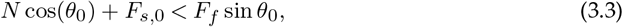

Equations 3.1, 3.2, and 3.3 can be solved numerically to find the maximum normal force before slipping, which can then be used to find the maximum storable spring force (*F*_*s*,0_) (SI Fig. 7,8). From this, we find that as *µ* increases, the *F*_*s*,0_ also increases due to the increased friction force preventing slip for a greater range of forces. This follows the general statement made previously that higher friction improves the stability of the system, which allows it to store more energy. On the other hand, as *θ*_0_ increases, the *F*_*s*,0_ decreases (making the system less stable) because the greater initial angle results in a larger horizontal component of the normal force and a smaller horizontal component of frictional force (SI Fig. 7,8). For the purposes of these simulations, we use *θ*_0_ = 1*°* and *n* = 0.85 [36].

We use these three additions to the base LaMSA model to develop a series of equations that can be solved for any system to determine the evolution of the system over time. First, we write a force balance on the load mass (*m*) as follows:

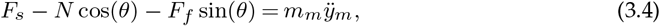

where *F*_*s*_ represents the spring force, *θ* represents the angle between the center of the latch and the point of contact between latch and load mass, and *mm* and *ÿm* represent the mass and acceleration of the load mass respectively. Similarly, we write a force balance on the latch:

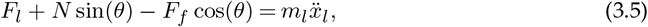

where *F*_*l*_ represents the force of the unlatching motor on the latch and *m*_*l*_ and 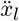 represent the mass and acceleration of the latch respectively. While the latch and load mass are in contact, they follow the circular curve of the latch, represented by:

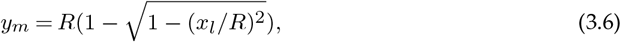

where *y*_*m*_ represents the y position of the load mass, *R* represents the radius of the latch, and *xl* represents the x position of the latch. We take the second derivative of this equation to yield the following equation linking the accelerations of the latch and load mass:

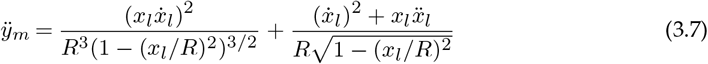

By combining Equations 3.1, 3.4, 3.5, and 3.7, we derive an ordinary differential equation relating the accelerations of the latch and load to the system properties (*R, µ, n, m*_*m*_, *m*_*l*_) and the forces acting on the latch and load mass at any given time (*Fs, Fl*).

We solve this equation using a 4,5 Runge-Kutta ODE solver(ode45) to determine the evolution of the model over time for as long as the latch remains in contact with the load mass. At *t <* 0, the spring is loaded by a motor to an initial spring force *F*_*s*,0_. At *t* = 0, the unlatching motor begins to exert an unlatching force on the latch and causes it to accelerate in the positive x direction. During this time, as the latch accelerates away, the load mass begins to accelerate, driven by releasing of the spring force (Fig. 5A.) The release of elastic energy as the spring returns to equilibrium causes the spring force acting on the load to decrease. Combined with the acceleration of the latch away from the load, the normal force between the load and the latch decreases (Fig. 5C). This eventually leads to a complete detachment of load mass and latch, which can bee seen when the normal force drops to zero (Fig. 5C). We define to this as the unlatching point and call the time taken to reach this point the unlatching time (*t*_*ul*_). After this unlatching, the load mass continues to accelerate as the remaining elastic energy in the spring is released. Once all elastic energy is released and the spring has returned to equilibrium, the load mass has reached the maximum velocity attainable. We define this point to be the take-off point and accordingly term this maximum velocity the take-off velocity (*v*_*TO*_). We assume the load mass detaches from the spring at the take-off point and travels with no external forces applied (Fig. 5B). In this model, *v*_*TO*_
is analogous to *ω*_*max*_ experimentally since *ω*_*max*_ represents the peak finger velocity after unlatching.

**Figure 5.**
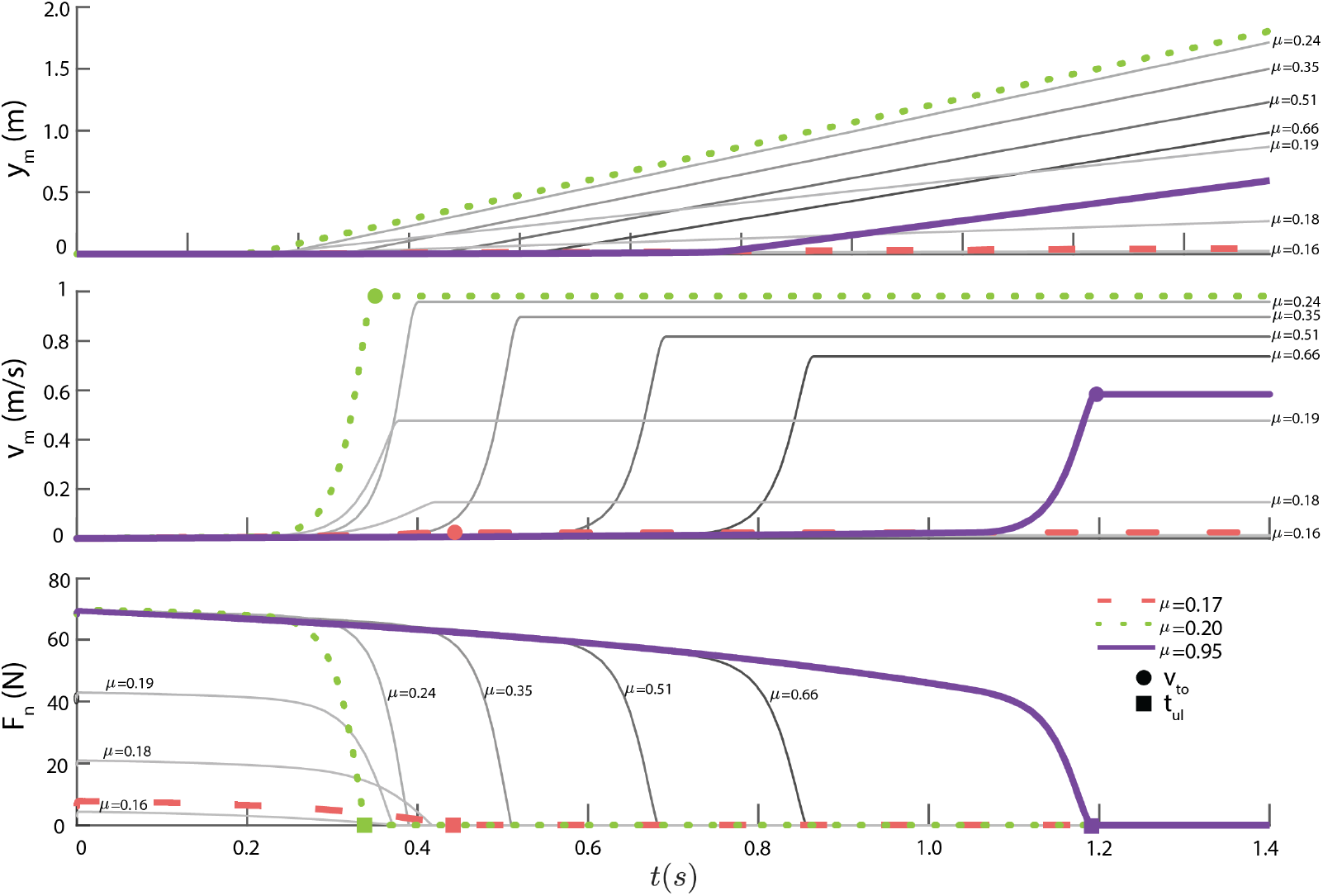
The model output is qualitatively similar to the kinematics of finger snapping experiments. **A**. The position of the load mass (*y*_*m*_) increases slowly until the system unlatches (occurring when *N* reaches 0). After this point, the load mass follows simple harmonic motion until it reaches its maximum velocity, at which point it takes off from the spring and continues at this velocity. **B**. The velocity of the load mass (*v*_*m*_) increases from zero until the system unlatches. The load mass follows simple harmonic motion until maximum velocity (*v*_*to*_) is reached, which is maintained after take off. The model shows an optimal *µ* of 0.20, as at *µ* lower or higher than this, the *v*_*to*_ decreases. **C**. The normal force acting on the load mass (*N*) begins at its maximum before decreasing to zero. When *N* = 0, the system has unlatched (*t*_*ul*_). The previously noted optimal *µ* produces the lowest *t*_*ul*_ while lower and higher *µ* leads to higher *t*_*ul*_.

By continuously tuning the frictional coefficient in the model, we find two important roles of friction. First, at low friction coefficients, the friction force between latch and load remains low, limiting the amount of force the loading motor can compress the spring to before slipping. As *µ* increases however, the friction force between latch and load grows, increasing the force that can be loaded before slipping and leading to an increase in the take-off velocity (Fig. 6A). At higher values of *µ*, the additional constraint of a maximum loading motor force limits the force loaded. Therefore at large values of *µ*, the stored force can no longer increase (Fig. 6A). Instead, increasing *µ* causes greater energy dissipation during unlatching which causes a longer unlatching time (Fig. 6B). This dissipation ultimately reduces the take-off velocity (Fig. 6C).

**Figure 6.**
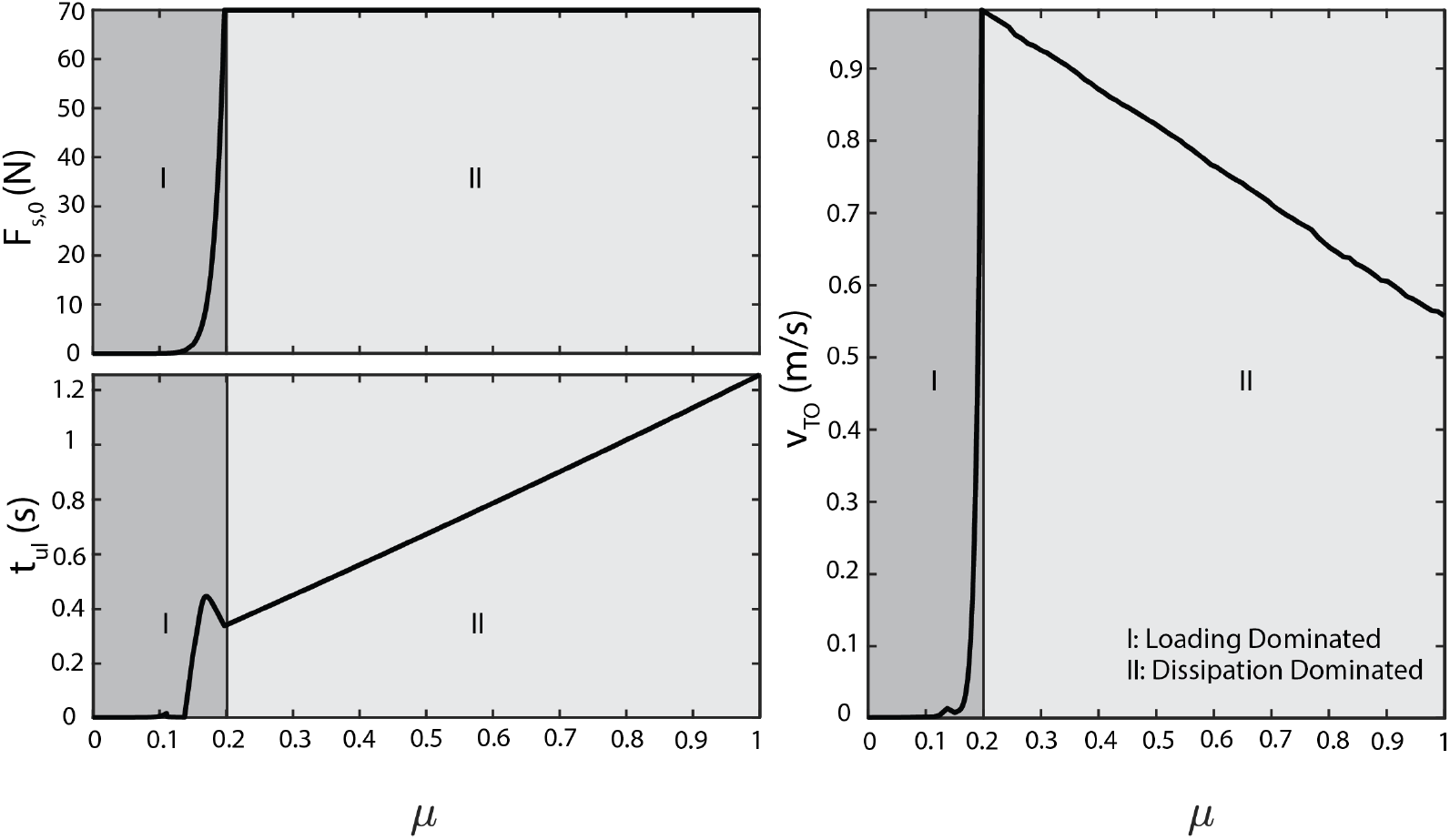
Increasing *µ* results in unique trends with respect to *v*_*to*_ and *t*_*ul*_. **A**. As *µ* increases, the loaded *F*_*s*,0_ increases until it reaches the limit of what the system can store. **B**. While in region 1, a loading dominated regime, *t*_*ul*_ decreases with *µ* due to the increase in *F*_*s*,0_. However, once in region 2, an dissipation dominated regime, *t*_*ul*_ increases with *µ* as more energy is converted to frictional energy. **C**. While in the loading dominated regime, *v*_*to*_ increases as *F*_*s*,0_ increases, providing greater stored spring energy that is converted to kinetic energy. However, the system then transitions to an dissipation dominated regime where *F*_*s*,0_ remains constant. This results in *v*_*to*_ decreasing with *µ* because more energy is lost to friction, as represented by the increase in *t*_*ul*_. Overall, this results in a peak in *v*_*to*_ occurring at the transition between the loading dominated and dissipation dominated regimes.

In summary, as *µ* increases from low values, more energy can be stored in the spring without slipping, increasing the *v*_*TO*_
. Thus, we term this region “loading dominated.” However, as the *µ* continues to increase, the maximum force achievable by the loading motor is reached. This limits the elastic energy stored in the spring while continuing to increase the energy dissipated by friction, as shown by the increasing *t*_*ul*_. The increased dissipation with no increase in energy storage leads to decreasing *v*_*TO*_
, and is thus named “dissipation dominated”. We note that these trends in *F*_*s*,0_, *t*_*ul*_, and *v*_*TO*_
qualitatively match what is seen experimentally in the finger snap experiments (Figs. 6 and 5).

Examining how the stored elastic energy and dissipated energy depend on friction reveals a general condition for when we might expect optimal velocities at intermediate friction in a LaMSA system. Considering both the stored potential energy in the spring 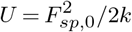 and the resulting maximum kinetic energy 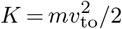, the total energy dissipated by the latch is the difference between these, namely, *Ed* = *U − K*. We expect an optimum in *v*to(*µ*) to occur when 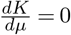, or in other words when

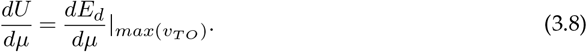

We validate this condition by determining the total dissipated energy and stored spring potential energy for varying values of *µ*. The results, shown in Figure 7, show that where condition 3.8 is true corresponds to the highest *v*_*TO*_
the system was able to achieve by varying *µ*. While operating within the loading dominated regime, the elastic potential energy, total dissipated energy, and maximum kinetic energy increase with frictional coefficient (Fig. 7A). However while in the dissipation dominated regime, the imposed biological maximum keeps the elastic potential energy constant but the total dissipated energy increases and the maximum kinetic energy decreases due to higher frictional coefficient (Fig. 7A). Since the elastic potential energy is constant while the total dissipated energy continues growing at the transition between the two regimes, 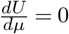 and 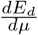 remains nonzero, becoming equal to each other at this transition (Fig. 7B). Therefore, Eq. 3.8 holds true for the *µ* with the maximum *v*_*TO*_
.

**Figure 7.**
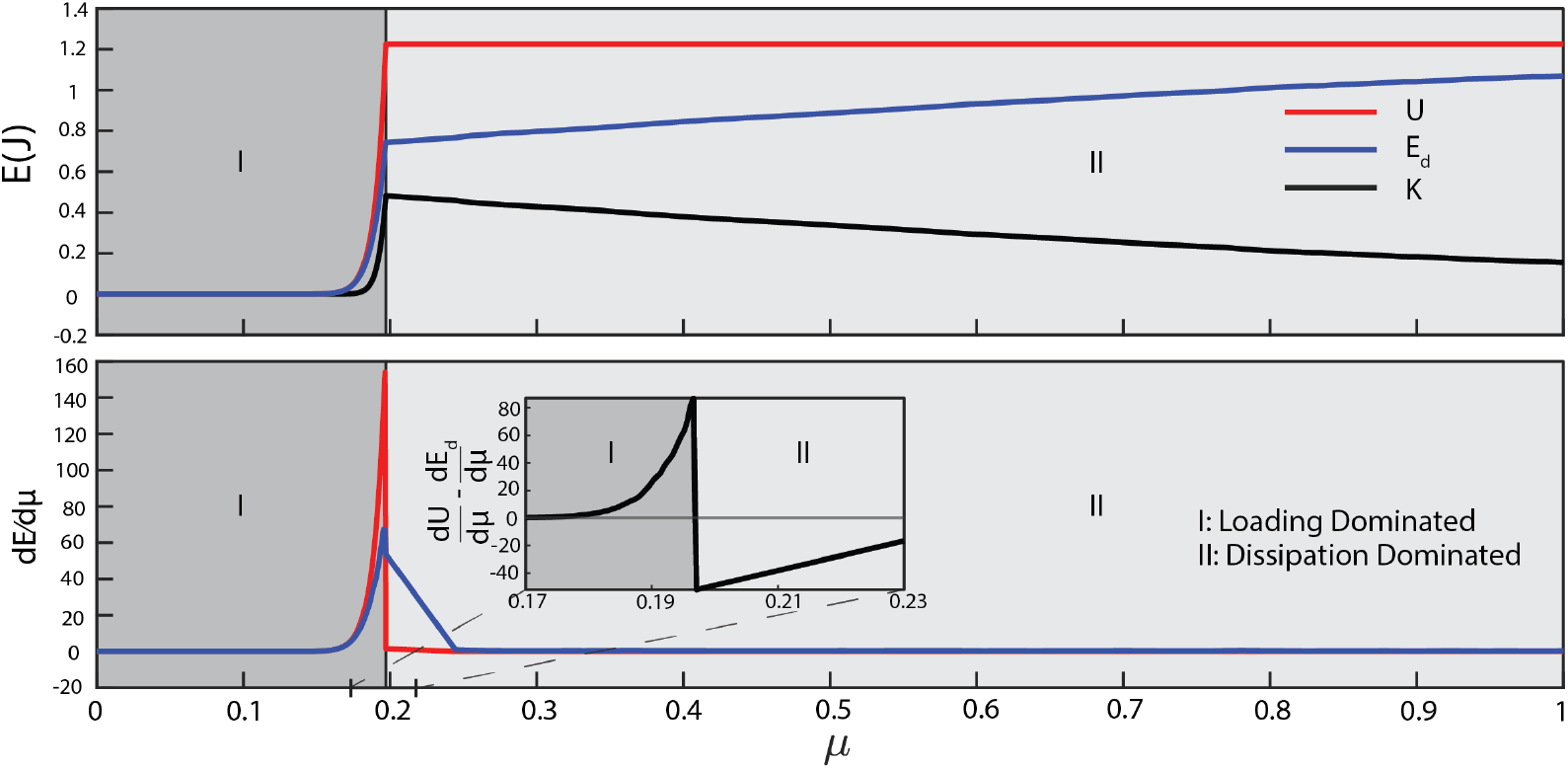
Peak *v*_*to*_ can be predicted by energetic analysis. **A**. Shown are the maximum kinetic energy achieved by the load *K* (black line), initial potential spring energy *U* (blue), and energy dissipated *E*_*d*_ (red) for each point *µ* from the soft body model. As *µ* increases, *U* increases until it reaches the maximum storage capacity of the system while *K*_*max*_ achieves a peak before decreasing. *E*_*d*_ consistently increases with increasing *µ*. **B**. We calculated the the derivatives of potential energy and dissipated energy with respect to *µ* (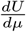 and 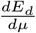) for the phenomenological model. It can be observed that the peak in *K* occurs at a *µ* where the difference in 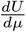 and 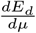 intersect, the point which marks the transition from a loading dominated regime to a dissipation dominated one.

This energy analysis reveals the physical reason that a frictional LaMSA system with instability (*θ*_0_ *>* 0) must have a peak in the take-off velocity: the dual role of friction in the LaMSA system. In the loading dominated regime, the latching capability of the friction plays a greater role than its dissipative role, allowing for greater energy storage and thus greater *v*_*TO*_
. However, this latching capability is limited by the capabilities of the spring and motor, eventually leading to the dissipative effects of friction overtaking the latching capability, which results in the eventual decrease in *v*_*TO*_
.

In summary, this model shows similar trends to those observed experimentally. In the finger snap system, we observe that *ω*_*max*_ has a peak with respect to *µ* and that this peak likely occurs from the interplay between dissipated energy (*∝ t*_*ul*_) and stored energy (*∝ F*_*max*_), both of which increase with *µ* (Fig. 3). In our model, we show that both the *F*_*s*,0_, analogous to *F*_*max*_, and *t*_*ul*_ of the modeled system increase with *µ* and that these trends result in a peak in *v*_*TO*_
, which is analogous to *ω*_*max*_ in the finger snap system (Fig. 6). Using this model, we are also able to elucidate how the interplay between stored and dissipated energy, hinted at by the experimental results, defines this peak *v*_*TO*_
(Fig. 7).

### (d) Capabilities and Limitations of this Model

We had two goals in creating this model. First, this model seeks to identify the key components of the finger snap that enable the ultra-fast motion we observe. Second, this model seeks to capture the importance of friction in a friction-based LaMSA system such as a finger snap. This model was not meant to comprehensively replicate the physical or mechanical properties of the features involved (muscles, skin friction, etc.)

The major advantage of this LaMSA model with soft body friction is the capability to qualitatively capture the trends relevant to the finger snap system. Although the constraints tested here are particular to the finger snap, we tested other sets of assumptions and show that they too are able to reflect experimental results (SI Fig. 4, 5, 6, 7). This shows that aside from being qualitatively reflective of trends, this modeling approach allows for a great degree of flexibility with the system being analyzed. This model is capable of accommodating multiple types of springs, loading motors, unlatching motors, and a limited variety of frictional models. This flexibility allows the model to be more broadly applicable and is one potentially useful way of exploring friction in other frictional and soft body LaMSA systems.

However, this model has several notable limitations. Although the main frictional model discussed here has been used in soft body friction modeling in finger pads, it typically is only applicable in forces of up to 1 N and in some cases is a poor model for soft body, viscoelastic friciton. Thus, this may not be the best model to use in capturing the nonlinearities of such systems [35, 37]. However, current models of skin friction as a result of deformation depend on the change in surface area of the finger pad [34, 38]. This model decouples the change in geometry of the compressible surface from the effect this has on friction using the empirical model described above. However, the change in geometry is likely a significant factor which this model is currently unable to support. Therefore, while the qualitative trends with respect to soft-body friction can be gleaned, this model is limited in accurate quantitative predictions and should only be used to observe and qualitatively understand the trends of a given LaMSA system.

## 4. Concluding Remarks

The finger snap is a fascinating LaMSA system. We have shown that the loading and unlatching are both heavily mediated by the latch friction, which makes this an excellent example of a friction based LaMSA system. With our model we explore this dual role of friction in the finger snap and how it gives rise to a maximum take off velocity *v*_*TO*_
with an optimum coefficient of friction *µ*. Through an energetic analysis this optimum *µ* can be identified for any generic system, opening this analysis to other friction based LaMSA systems.

Our model suggests that for certain latch-spring systems, friction can play two competing roles: aiding in the loading of the spring and hindering the unlatching of the latch and load mass. This unique trade off in energy due to the dual role of friction has been previously explored in granular flow systems [39], but can now also be applied to ultrafast snapping LaMSA systems as well. For example, certain systems such as the Panamanian termite soldier (*Termes* panamaensis) [8] or the Dracula ant (*Myrmoteras* formicinae) [10] can generate a high speed motion using a similar snap movement that may heavily rely upon friction. Using the framework established here, these LaMSA systems can be analyzed with a greater understanding of the underlying frictional mechanics. Moreover, the optimization analysis discussed here shows that the friction can be used as a method of fine-tuning the motion of a system, which can inform improve prosthesis designs [40, 41, 42]. For soft robotics, achieving realistic models of friction dynamics has been identified as an avenue for improving the manipulative capabilities of robotic systems [43, 44]. Thus, our work has potential to contribute to a growing body of bio-inspired latches where robust modulating ability of frictional latches can enhance and pave new functionalities in soft robots [33, 45, 46].

## 5. Code and Data Availability

All code and datasets presented in this paper are available at this link: https://github.com/bhamla-lab/FingerSnap_2021

## Acknowledgment

We thank members of the BhamlaLab for useful discussions and feedback. We thank two members of the BhamlaLab for volunteering to participate in the finger snap experiments. In particular we thank Harry Tuazon for valuable assistance in development of experiments for measurement of frictional coefficients. R.A. acknowledges funding support from Georgia Tech’s Presidential Undergraduate Research Award (PURA). M.S.B acknowledges funding support from NSF CAREER award no. 1941933. M.I. acknowledges funding support from the NSF for this work under grant no. 2019371.

## 6. Author Contributions

RA performed the experiments, participated in data analysis, performed the modeling, and wrote the manuscript. EJC performed the experiments, participated in data analysis, and helped draft and critically revise the manuscript. MI performed the modeling and helped draft and critically revise the manuscript. MSB conceived of this study, participated in data analysis, performed the modeling, and helped draft and critically revise the manuscript. All authors gave final approval for publication and agree to be held accountable for the work performed therein.

