## Supplementary material for "The ultrafast snap of a finger is mediated by skin friction": SI pdf

ROYAL SOCIETY OPEN SCIENCE

rsos.royalsocietypublishing.org

Research

Article submitted to journal

**Subject Areas:**
Biomechanics, Biophysics

**Keywords:**
Finger snap, Latch Mediated Spring Actuation (LaMSA), Ultra-fast motion

**Author for correspondence:**
Saad Bhamla


### The ultrafast snap of a finger is mediated by skin friction

Raghav Acharya[1], Elio J. Challita[2], Mark Ilton[3] and M. Saad Bhamla[1]

[1] Chemical and Biomolecular Engineering, Georgia Institute of Technology, Atlanta, GA 30311, USA
[2] George W. Woodruff School of Mechanical Engineering, Georgia Tech, Atlanta, GA,30311, USA
[3] Department of Physics, Harvey Mudd College, Claremont CA, USA, 91711

#### Contents

**1 Contents of SI Movies** 2

**2 SI Tables** 2

**3 SI figures** 3
(a) Alternate ways to model finger snaps . . . 3

© 2014 The Authors. Published by the Royal Society under the terms of the Creative Commons Attribution License http://creativecommons.org/licenses/by/4.0/, which permits unrestricted use, provided the original author and source are credited.

THE ROYAL SOCIETY
PUBLISHING

rsos.royalsocietypublishing.org R. Soc. open sci. 000000

### 1. Contents of SI Movies

(i) **SI Movie S1: Normal Motion of a Fingersnap** Recording of unaltered finger snap from side at 4082 fps, played at 60 fps and from front at 4082 fps, played at 3000 fps (35 times speed).

(ii) **SI Movie S2: Comparison between Fingersnaps covered by Nitrile and Lubricated Nitrile** Recordings of nitrile covered finger snap and lubricated finger snap at 4082 fps, played at 60 fps (17.5 times speed).

(iii) **SI Movie S3: Comparison between Fingersnaps covered by Nitrile and Latex Rubber** Recordings of nitrile covered finger snap and Latex Rubber covered finger snap at 4082 fps, played at 60 fps (17.5 times speed).

(iv) **SI Movie S4: Comparison between Nitrile covered Fingersnaps and Latchless motor driven finger motion** Recordings of nitrile covered finger snap and latchless finger motion at 4082 fps, played at 60 fps (17.5 times speed).

(v) **SI Movie S5: Comparison between Nitrile covered Fingersnaps and Nitrile covered Thimble** Recordings of nitrile covered finger snap and nitrile covered thimble finger snap at 4082 fps, played at 40 fps (17.5 fps).

### 2. SI Tables

| Snapper | p |
|---|---|
| 1 | $1.45 \times 10^{-6}$ |
| 2 | $1.04 \times 10^{-6}$ |
| 3 | $3.45 \times 10^{-6}$ |

**Table 1.** ANOVA p values for maximum angular velocities observed for each snapper between snaps covered in different materials. Each snapper shows significant differences in the means of each condition tested (nitrile, lubricated nitrile, latex rubber).

| Snapping condition | p |
|---|---|
| Nitrile | 0.752 |
| Lubricated Nitrile | 0.982 |
| Latex Rubber | 0.922 |

**Table 2.** ANOVA p values for maximum angular velocities observed for each snap condition between snappers. For each condition tested (nitrile, lubricated nitrile, latex rubber), each snapper exhibited similar maximum angular velocities.

| Feature | p |
|---|---|
| $F_{max}$ | 0.0124 |
| $t_{ul}$ | 0.493 |

**Table 3.** Pairwise t-tests p values for Force Dynamic features of "strong" finger snaps and "weak" finger snaps

### 3. SI figures

#### (a) Alternate ways to model finger snaps

While we show that the best way to model the finger snap is with a soft body model of friction, there are other ways to reveal a similar trend. In order to do so, the main trend that the initial spring force increases with $\mu$ must be maintained. We do so by imposing an artificial loading constraint which follows:

$$F_{s,0} = \alpha \times \mu + F_0 \tag{3.1}$$

Where $\alpha$ represents the relationship between $F_{s,0}$ and $F_0$ represents an initial spring force that is loaded at $\mu$=0. The results of these models are shown in Figs (9,10,11,12).

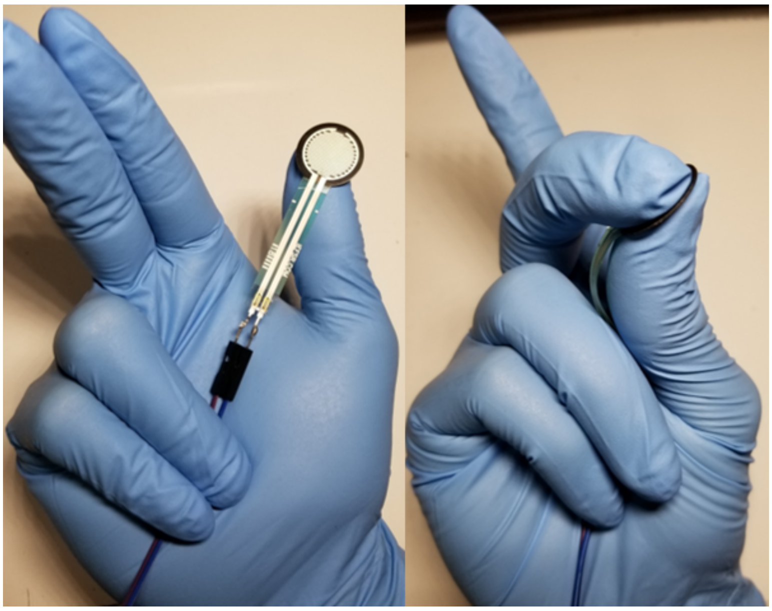

**Figure 1. Positioning of Force Sensor in Force Dynamics Experiments** An FSR interlink 402 was placed between thumb and middle finger to obtain force data from the snap. This FSR was connected to a voltage divider and the output voltage is fed to the analog input of an Arduino UNO (https://www.sparkfun.com/products/9375?_ga=2.37027559.611372374.1629255380-422841328.1629255380). A custom Arduino script converts the voltage values to their corresponding force values as per the established documentation. This was tested by placing items of known mass (3 kg to 10 kg) on the FSR and ensuring the calculated force corresponded to the force of gravity exerted upon the block.

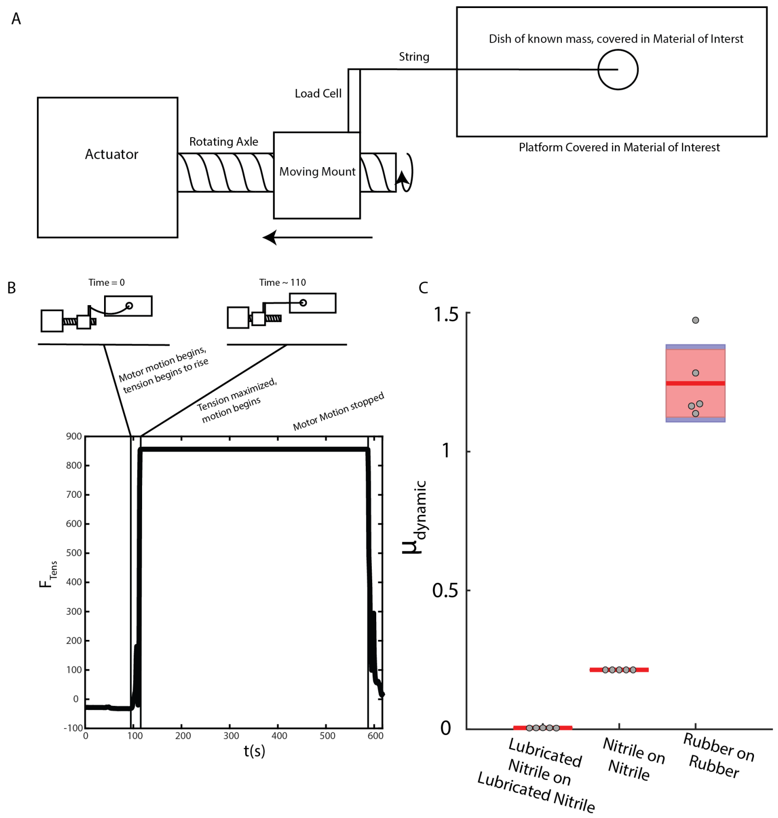


**Figure 2. Summary of Experiments to measure friction coefficient ($\mu$) A.** A schematic of the experiment performed to determine the friction coefficient between any two materials. One material is affixed to the dish and the other to the platform. The dish is filled with some known weight $m_{dish}$ and pulled by the linear actuator. A load cell connected with a rigid line to the dish is used to measure the horizontal force resisting the movement. **B.** An example force profile measured by the load cell over the course of one trial. The measured force is initially close to 0 as there is little force on the load cell. Once the motor begins to move, the moving mount retracts, increasing the tension in the string until the dish begins to move. During motion, which is at a constant velocity, the measured force is constant ($F_{peak}$. This force is used in combination with the known weight to calculate the friction coefficient of the materials used through the following equation: $\mu = \frac{F_{peak}}{m_{dish}g}$. The temporal resolution of the load cell used is not sufficient to consistently or accurately determine the static friction coefficient, but we are able to determine the dynamic friction coefficient. **C.** Summary of calculated $\mu_{dynamic}$ for each of the three combinations of materials. As expected, the lubricated nitrile on lubricated nitrile exhibited the lowest friction coefficients while latex rubber on latex rubber exhibited the greatest friction coefficients.

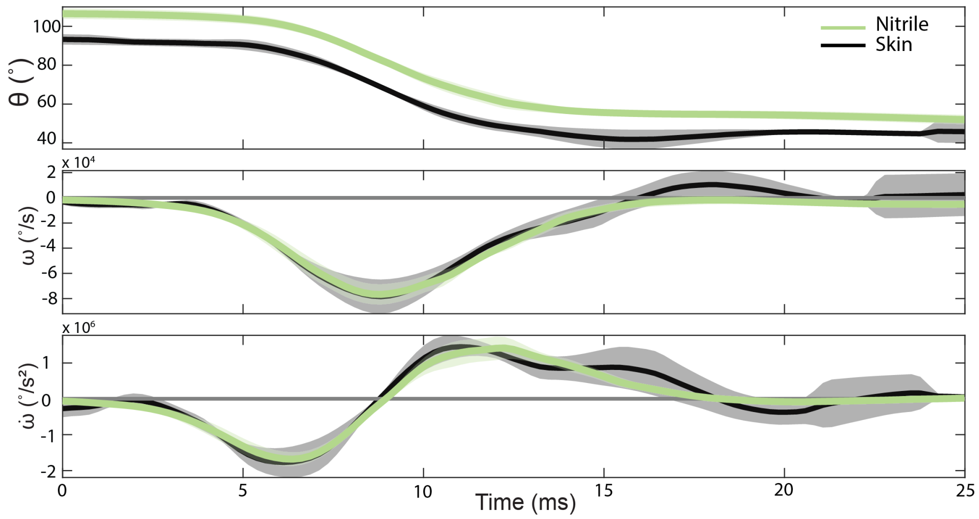


**Figure 3. Nitrile covering the fingers do not change snap kinematics.** Data from high speed videos taken at 4082 fps (N=5 each) and smoothed using 17 point ($\sim 5\%$ smooth) Savitsky-Golay filter. The angular displacement, velocities, and accelerations are identical. Nitrile is a proper substitute of dry skin as it replicates the kinematics and overall general trends of dry skin. This allows us to make controllable modifications to the $\mu$ and eliminates variability in surface due to sweat and humidity.

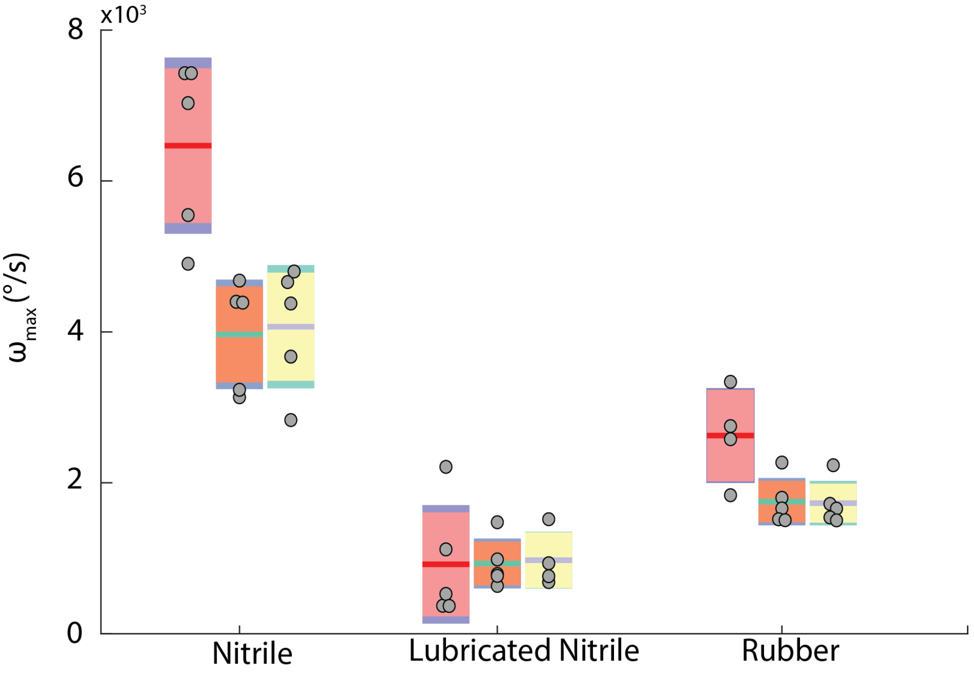


**Figure 4. The maximum angular velocity of three different individual snappers is preserved.** Although for certain conditions, different snappers exhibited significant differences in snap velocities, for each individual snapper, the trend of the nitrile snap having the highest $\omega_{max}$ compared to other conditions held true. ANOVA tests were performed to analyze the variance of means for each condition and for each snapper and the results of these tests can be found in Tables 1 and 2.

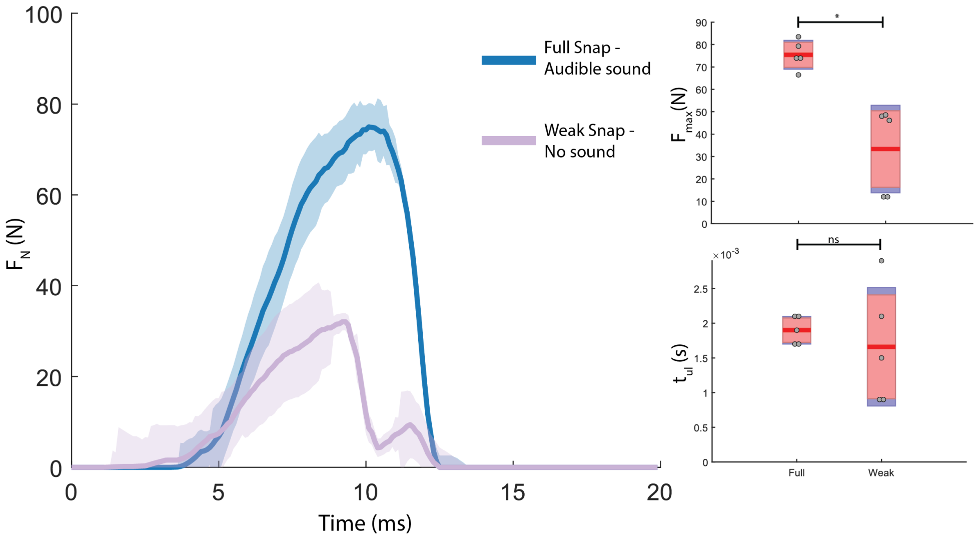


**Figure 5. The force dynamics of a "strong" snap with an audible sound are different from a "weak" snap with no sound. A.** The force profiles reveal that there are notable differences in the force profile between a "strong" snap, defined as one which produces an audible "pop" sound, and a "weak" snap, defined as one which produces no such sound. **B.** Weak snaps have a lower maximum force when compared to strong snaps. **C.** Weak and strong snaps have similar unlatch times. One-way t-tests were performed, and the results of these tests can be found in table 3

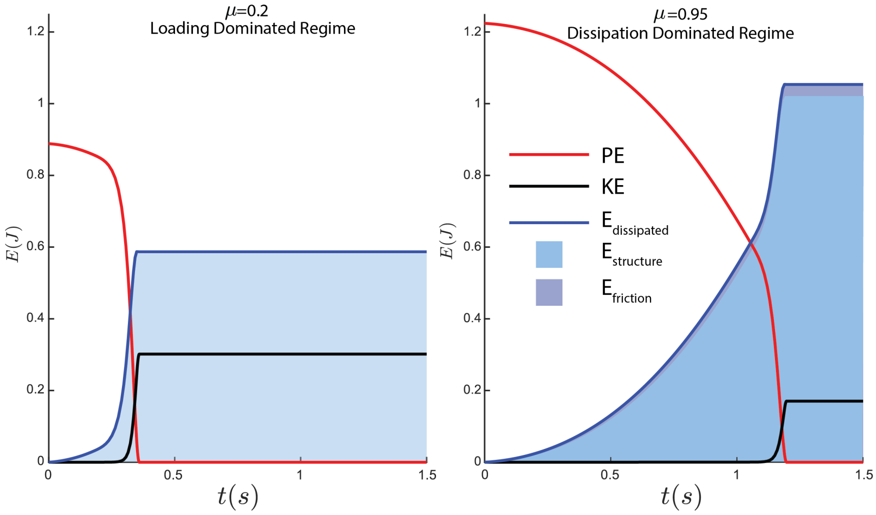


**Figure 6. Increasing friction coefficient results in a greater dissipation of energy due to friction in a compressible model.** Here we show the transformation of stored potential energy through time for the compressible model described in the Results and Discussion section. Initially, all energy is stored in the spring as potential energy. The energy is converted to kinetic energy or is dissipated as the system unlatches. Energy is dissipated either due to latch opposing motion ($E_{structure}$) or friction ($E_{friction}$). As the $\mu$ increases, several notable changes are observed. A greater quantity of the stored energy is dissipated while less is converted to kinetic energy. Within the dissipated energy, more energy is dissipated due to friction as $\mu$ increases. Additionally, the transfer of potential energy takes longer as $\mu$ increases.

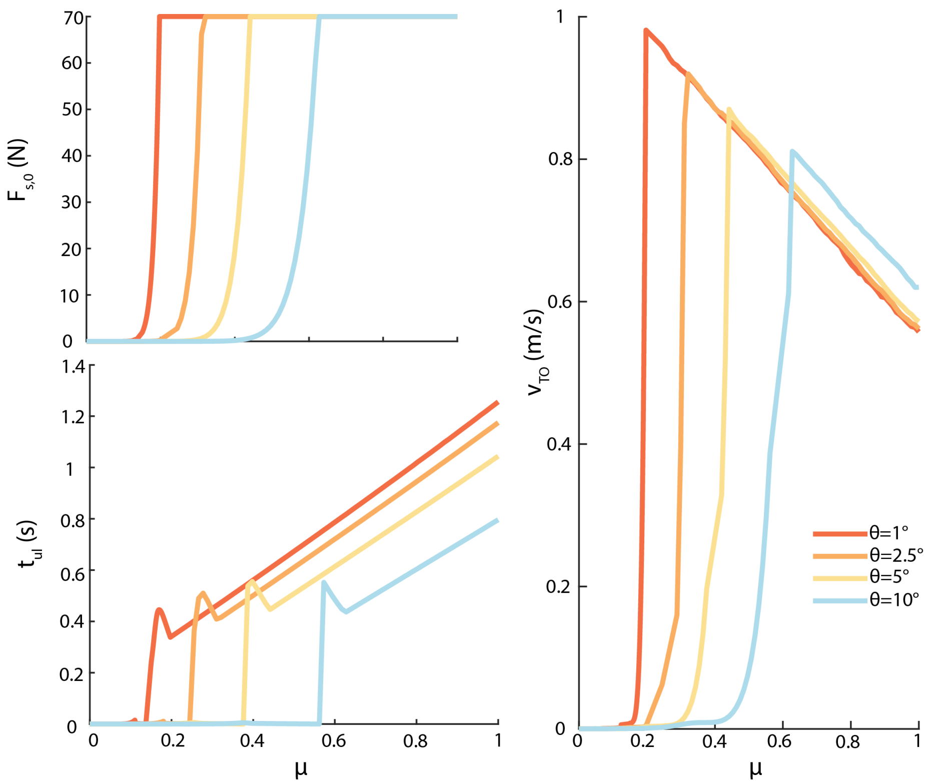


**Figure 7. Increasing $\theta_0$ in a compressible model leads to systems with less stability. A.** The increasing $\theta_0$ causes the $F_{s,0}$ to increase slower. This occurs because the greater $\theta_0$ leads to a greater horizontal component of normal force and smaller horizontal component of the frictional force, reducing the $F_{s,0}$ with $\theta_0$ for a given $\mu$. These results can also be interpreted as increasing $\theta_0$ resulting in greater instability for the system, as in more unstable systems (higher $\theta_0$), the system is unable to store as much force. **B.** Increasing $\theta_0$ results in decreased $t_{ul}$ at most $\mu$. This decrease is due to the fact that by starting at a higher $\theta_0$, there is less distance on the latch that the load needs to travel which allows unlatching to occur faster. **C.** Greater $\theta_0$ results in systems which reach a peak $v_{TO}$ at higher friction coefficients. This occurs because the greater $\theta_0$ increases horizontal normal force components while decreasing horizontal friction force components, reducing $F_{s,0}$ with $\theta_0$ for a given $\mu$. Conceptually, this can be understood as systems with higher values of $\theta_0$ being inherently less stable and requiring higher friction coefficients to be able to support spring loading at the maximum capability of the loading motor. As a result of this required increase in $\mu$, at higher values of $\theta_0$, the peak $v_{TO}$ is greater as less energy is dissipated at these lower values of $\mu$.

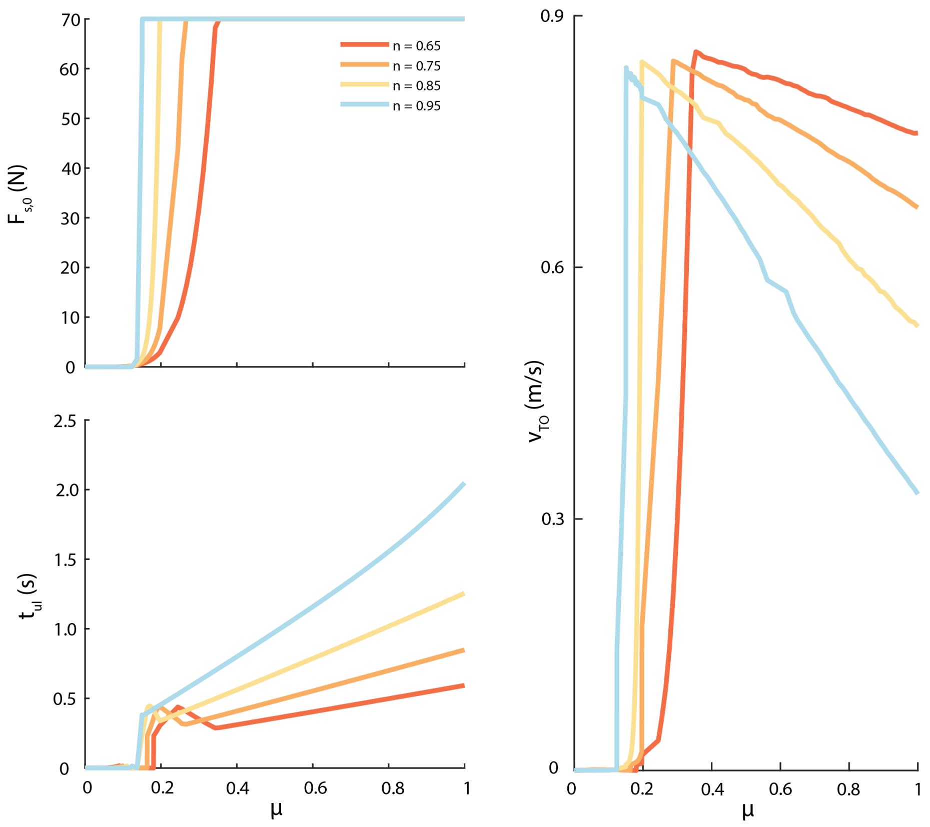


**Figure 8. Increasing compressibility by decreasing $n$ in a compressible model allows systems to achieve higher peak velocities. A.** As $n$ decreases (compressibility increases), the increase in $F_{s,0}$ occurs at reduced rates. This trend occurs because the increased compressibility reduces the friction force the system can generate for a given $\mu$, resulting in reduced energy storage capability. **B.** Increasing $n$ (reducing compressibility) results in greater $t_{ul}$ for most $\mu$. Although the greater $n$ results in a more rapid increase in $F_{s,0}$, once all systems reach the maximum $F_{s,0}$, systems with lower $n$ experience less friction, leading to lower $t_{ul}$ **C.** Lower $n$ (greater compressibility) leads to greater peak $v_{TO}$ values which occur at larger $\mu$. This apparent contradiction occurs because of the nonlinear nature of friction in a compressible system. For systems with greater compressibility (lower $n$), for a given $\mu$, the friction force is decreased. As a result, although the peak $v_{TO}$ values occur at greater values of $\mu$ due to slower $F_{s,0}$ increases, the friction is lower at these values, resulting in higher peak $v_{TO}$ values.

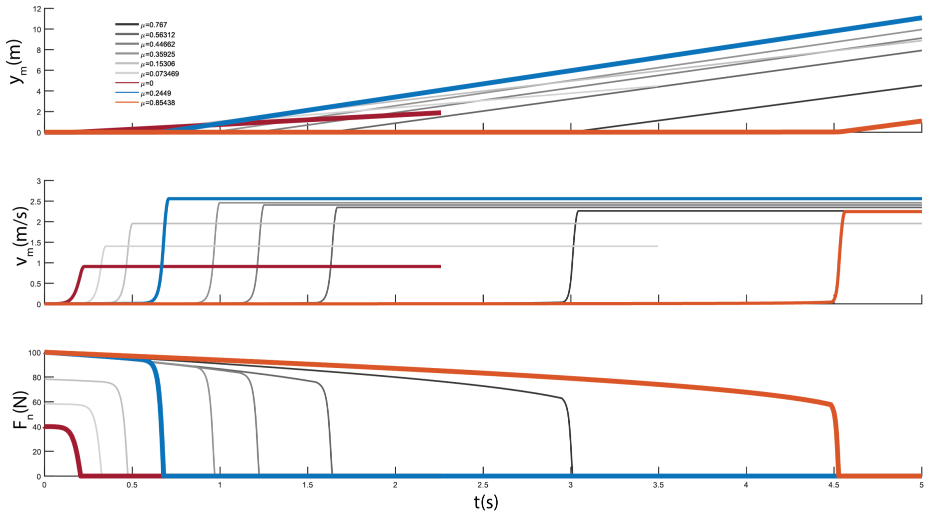


**Figure 9. A phenomenological model can be used to relate trends to the finger snap system. A.** The position of the load mass $(y_m)$ increases slowly until the system unlatches (occurring when $N$ reaches 0). After this point, the load mass follows simple harmonic motion until it reaches its maximum velocity, at which point it takes off from the spring and continues at this velocity. **B.** The velocity of the load mass $(v_m)$ increases from zero until the system unlatches. The load mass follows simple harmonic motion until maximum velocity $(v_{to})$ is reached, which is maintained after take off. The model shows an optimal $\mu$ of $0.08$, as at $\mu$ lower or higher than this, the $v_{to}$ decreases. **C.** The normal force acting on the load mass $(N)$ begins at its maximum before decreasing to zero. When $N = 0$, the system has unlatched $(t_{ul})$. The previously noted optimal $\mu$ produces the lowest $t_{ul}$ while lower and higher $\mu$ leads to higher $t_{ul}$.

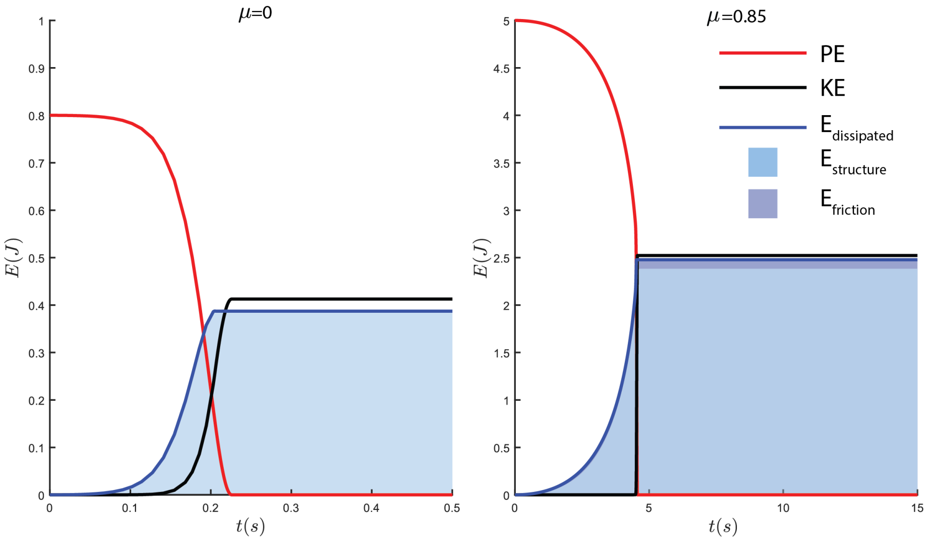


**Figure 10. Increased friction coefficients in a model with phenomenological loading results in greater dissipation of energy but may result in greater kinetic energy..** The transformation of energy through time of the system is shown. Initially, all energy is stored as potential spring energy, which is either converted to kinetic energy or dissipated as the system unlatches. The dissipated energy is lost due to either opposition of the latch or due to friction. Some energy is always lost due to the structure. As the coefficient of static friction increases from $\mu_s$ = 0 (**A**) to $\mu_s$ = 1 (**B**), the energy lost due to friction increases. However at the same time, as $\mu$ increases, the $PE$ also increases due to the imposed phenomenological loading. This increase allows the $KE$ to be greater at this higher $\mu$ than at the lower $\mu$, even though more energy is dissipated to the structure or to friction.

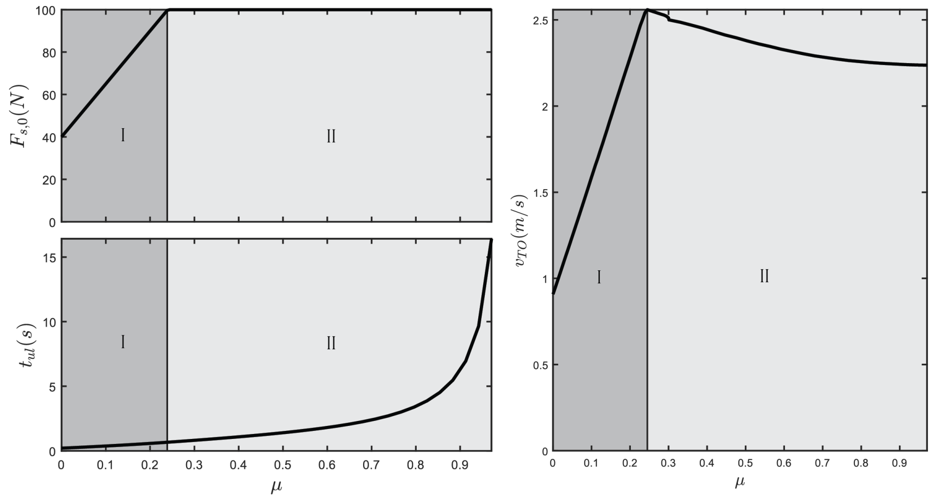


**Figure 11. Increasing $\mu$ results in unique trends with respect to $v_{to}$ and $t_{ul}$ when phenomenological loading is applied. A.** As $\mu$ increases, the loaded $F_{sp,0}$ increases until it reaches the limit of what the system can store following the phenomenological loading constraint applied to the system. **B.** While in region 1, a loading dominated regime, $t_{ul}$ decreases with $\mu$ due to the increase in $F_{sp,0}$. However, once in region 2, an unlatching dominated regime, $t_{ul}$ increases with $\mu$ as more energy is converted to frictional energy. **C.** While in the loading dominated regime, $v_{to}$ increases as $F_{sp,0}$ increases, providing greater stored spring energy that is converted to kinetic energy. However, the system then transitions to an unlatching dominated regime where $F_{sp,0}$ remains constant. This results in $v_{to}$ decreasing with $\mu$ because more energy is lost to friction, as represented by the increase in $t_{ul}$. This overall results in a peak in $v_{to}$ occurring at the transition between the loading dominated and unlatching dominated regimes.

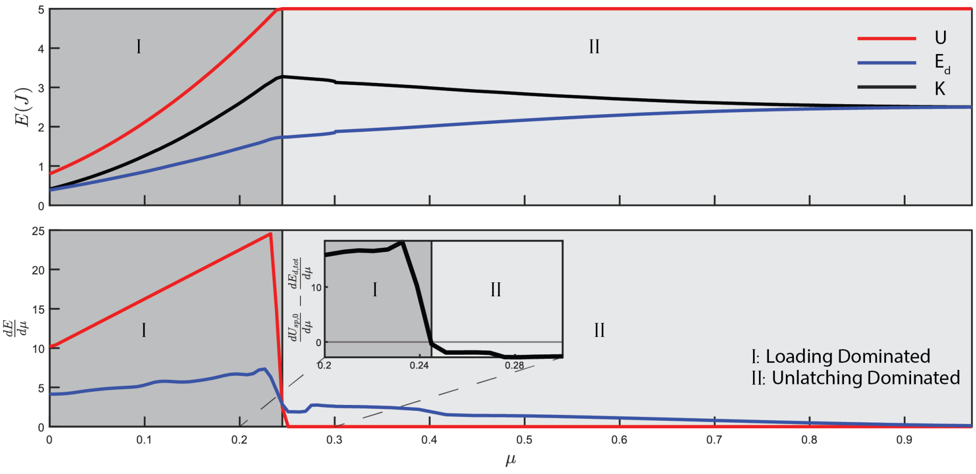


**Figure 12. Peak $v_{to}$ can be predicted by energetic analysis for a phenomenological loading model. A.** Shown are the maximum kinetic energy achieved by the load $K$ (black line), initial potential spring energy $U$ (blue), and energy dissipated $E_d$ (red) for each point $\mu$ from the phenomenological model. As $\mu$ increases $U$ increases until it reaches the maximum storage capacity of the system while $K_{max}$ achieves a peak before decreasing. $E_d$ consistently increases with increasing $\mu$. **B.** We calculated the the derivatives of potential energy and dissipated energy with respect to $\mu$ ($\frac{\partial U}{\partial \mu}$ and $\frac{\partial E_d}{\partial \mu}$) for the phenomenological model. It can be observed that the peak in $K$ occurs at a $\mu$ where the difference in $\frac{\partial U}{\partial \mu}$ and $\frac{\partial E_d}{\partial \mu}$ intersect, the point which marks the transition from a loading dominated regime to an unlatching dominated one.
